# Cortical neuronal assemblies coordinate with EEG microstate dynamics during resting wakefulness

**DOI:** 10.1101/2022.11.15.516587

**Authors:** Richard Boyce, Robin F Dard, Rosa Cossart

## Abstract

The disruption of cortical assembly activity has been associated with anesthesia-induced loss of consciousness. However, information about the relationship between cortical assembly activity and the variations in consciousness associated with natural vigilance states is currently lacking. To address this, we performed vigilance state-specific clustering analysis on 2-photon calcium imaging data from sensorimotor cortex in combination with global EEG microstate analysis derived from multi-EEG signals obtained from widespread cortical locations. Our analysis revealed no difference in the structure of assembly activity during quiet wakefulness (QW), NREMs, or REMs, despite the latter two vigilance states being associated with significantly reduced levels of consciousness relative to QW. However, we found a significant coordination between global EEG microstate dynamics and local cortical assembly activity during periods of QW, but not sleep. These results suggest that the coordination of cortical assembly activity with global brain dynamics could be a key factor of sustained conscious experience.

## Introduction

Neural assemblies, groups of neurons which coactivate together, have long been suggested to play an important role in cognition (Lorente de Nó, 1933; Hebb, 1949; Hopfield, 1982; Harris, 2005; Buzsaki, 2010). Several decades ago, the presence and proposed importance of assemblies in brain function began to find more experimental support as a result of advances in electrode recording techniques that enabled data to be obtained from large numbers of neurons spread over relatively localized brain regions (Carillo-Reid and Yuste, 2020). Such experiments were able to detect statistical features indicative of the presence of assembly organization (Wilson and McNaughton, 1993; Harris et al., 2003; Truccolo et al., 2010). More recently, the development of genetically encoded calcium indicators and optogenetic techniques in combination with 2-photon imaging platforms has allowed for the observation and manipulation of cortical neurons with a more optimal spatial resolution. Experiments completed as a result have enabled the direct visualization of localized microcircuit-level cortical assembly activity in mice *in vivo* (Miller et al., 2014), confirmed long-theorized properties attributed to microcircuit-level cortical assemblies such as pattern completion (Carillo-Reid et al., 2016), and demonstrated their importance in various cortical regions to associated learning behaviors (Carillo-Reid et al., 2019; Marshel et al., 2019).

The above results have cumulatively helped to establish local microcircuit-level cortical assemblies as critical units involved in cognitive tasks such as learning. Despite this, the more general question of their relationship with consciousness remains poorly understood. Using 2-photon imaging of layer 2/3 sensorimotor cortex in a head-fixed mouse model in addition to data obtained from high-density arrays implanted in humans, a recent study was able to gain valuable insight into local microcircuit-level cortical assembly dynamics in the awake versus anesthetized unconscious state (Wenzel et al., 2019). The authors reported that the onset of general anesthesia was associated with a reversible reduction and fragmenting of assemblies suggesting their potential role in maintaining consciousness and, more generally, supporting the hypothesis that cortical assemblies are unitary components of cognition (Hebb, 1949; Hopfield, 1982). However, there is currently a lack of information about local microcircuit-level cortical assembly dynamics across the full range of natural vigilance states, including sleep (Non-REM sleep (NREMs) and REM sleep (REMs)) when consciousness is significantly reduced relative to waking behaviors (Nir et al., 2013). If intact local microcircuit-level cortical assembly activity is a key component of consciousness, then this structured activity might also be expected to be altered to some extent during sleep.

To help improve our understanding of the potential role of local microcircuit-level cortical assemblies in consciousness, we have performed 2-photon calcium imaging of the sensorimotor cortex in combination with multi-EEG recordings across the full sleep-wake cycle in fully habituated head-fixed mice. Our initial aim was to compare and contrast the characteristics of local microcircuit-level assemblies occurring during each vigilance state in order to gain greater insight into their potential role in conscious experience.

## Results

### Imaging sensorimotor cortex across the full sleep-wake cycle

To facilitate stable long-term imaging of cortical neural activity, we injected an adeno-associated viral vector encoding the calcium indicator GCaMP6s driven under the hSyn promoter into the left ventricle of neonatal mice. The resulting brain-wide expression of GCaMP6s was stable well into adulthood (at least 6 months post-injection) (Fig. 1A; Fig. 1B, top right), as previously reported (Kim et al., 2014; Dard et al., 2022).

**Fig. 1.**
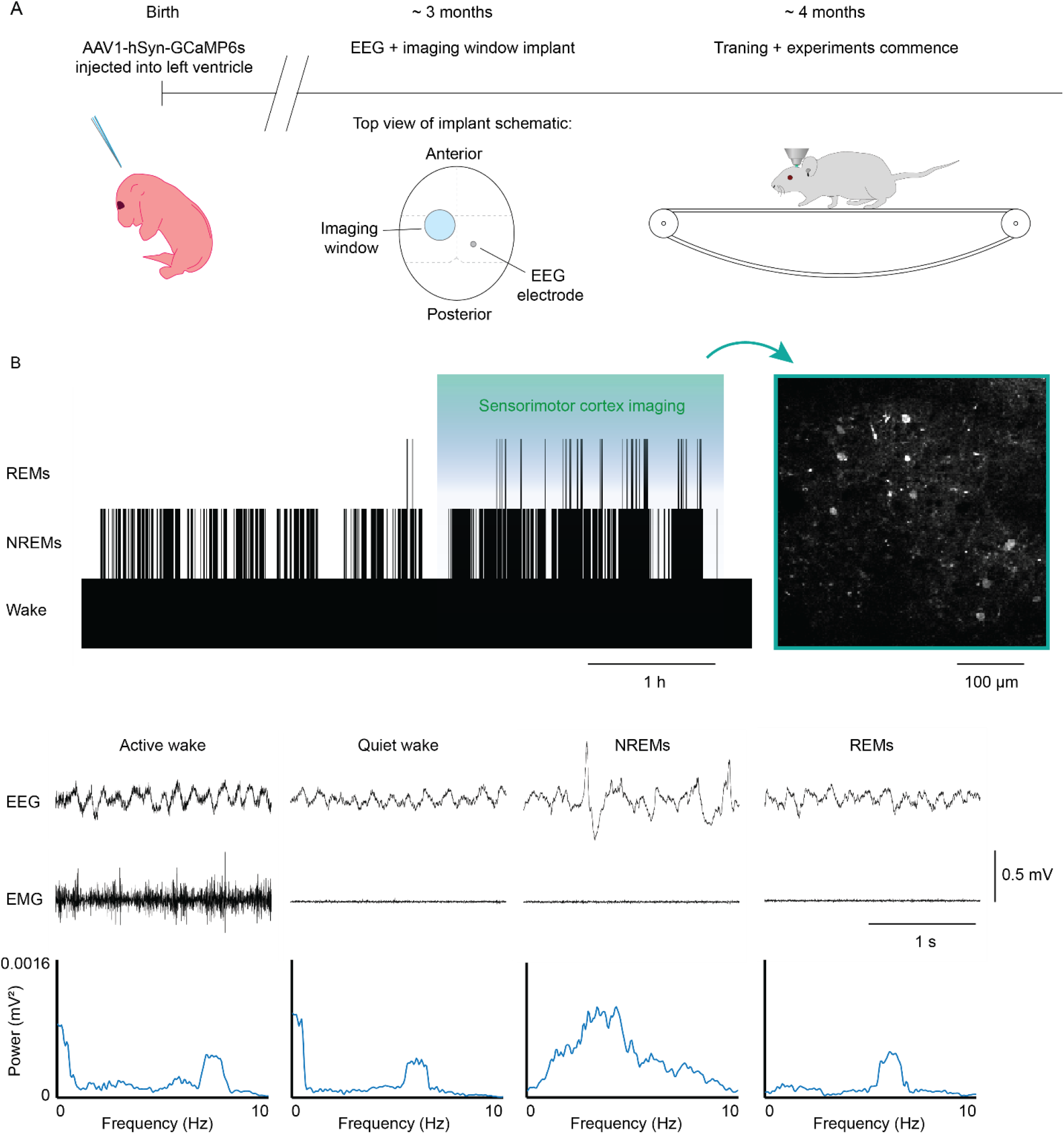
Imaging sensorimotor cortex across the full sleep-wake cycle. (**A**) Experimental timeline. Left. An adeno-associated viral vector encoding the calcium indicator GCaMP6s driven under the hSyn promoter was injected into the left ventricle of neonatal mice. Center. ~ 3 months later, mice were implanted with an imaging window and EEG and EMG electrodes used to assess sleep activity. Right. Following recovery from surgery, mice were trained to sleep on a non-motorized treadmill in order to image sensorimotor cortex across the full sleep-wake cycle. (**B**) Top. Representative hypnogram demonstrating the general imaging procedure in fully habituated mice. GCaMP6s expression resulting from P0 injection in neonates was widespread and stable long-term throughout the cortex allowing for high-quality imaging data acquisition from sensorimotor cortex months after the initial injection. To limit data amounts and minimize photobleaching, imaging data acquisition was completed during the second half of each experimental session when sleep activity was highest. Middle. Example EEG and EMG traces from major vigilance states analyzed in this study. Bottom. Spectral analysis derived from 5 s time epochs centered around the example EEG traces above. Spectral analysis of EEG signals in conjunction with EMG activity and behavioral observation was used to determine vigilance states throughout the entire course of experiments (see supplementary methods for details).

We next habituated previously-injected adult mice to sleep on a head-fixed non-motorized treadmill setup (Fig. 1A Fig. 1B, top). The habituation procedure took approximately 2 weeks to complete and was guided primarily through daily analysis of electrophysiological recordings derived from implanted EEG and EMG electrodes (see supplementary methods for details). Upon completion of habituation, all mice consistently exhibited a full sleep-wake cycle composed of periods of wakefulness, NREMs, and REMs (Fig. 1B). Following habituation, layer 2/3 of the sensorimotor cortex was continuously imaged through a window placed on the overlying dura in head-fixed mice in order to determine the characteristics of cortical assemblies across different vigilance states.

### Detection of vigilance state-specific local microcircuit-level assembly activity in sensorimotor cortex

Calcium trace-derived activity raster plots for a total of 1290 neurons from 6 mice (average = 215 ± 34 neurons per mouse) were extracted from the imaged movies (Fig. 2A). For use in all subsequent analysis, we isolated equal amounts of data for each of 4 vigilance states: active wakefulness (AW), quiet wakefulness (QW), NREMs (N), and REMs (R). Our first analysis focused on the activity of individual neurons. The proportion of cells active during periods of AW was significantly higher than for all other states, with the majority of all cells being active for at least 1 frame during the analysis period (AW = 84.0 ± 3.0 %; QW = 36.4 ± 10.3 %; N = 41.5 ± 11.7 %; R = 43.8 ± 11.5 %). Similarly, basic activity rates, measured as the number of active frames per second, were generally higher during periods of AW relative to all other states, with a significant difference in mean activity rate at the group level being found between AW and QW (Fig. 2C, i-ii). Neither of the above measures were significantly different between periods of QW, N, and R.

**Fig. 2.**
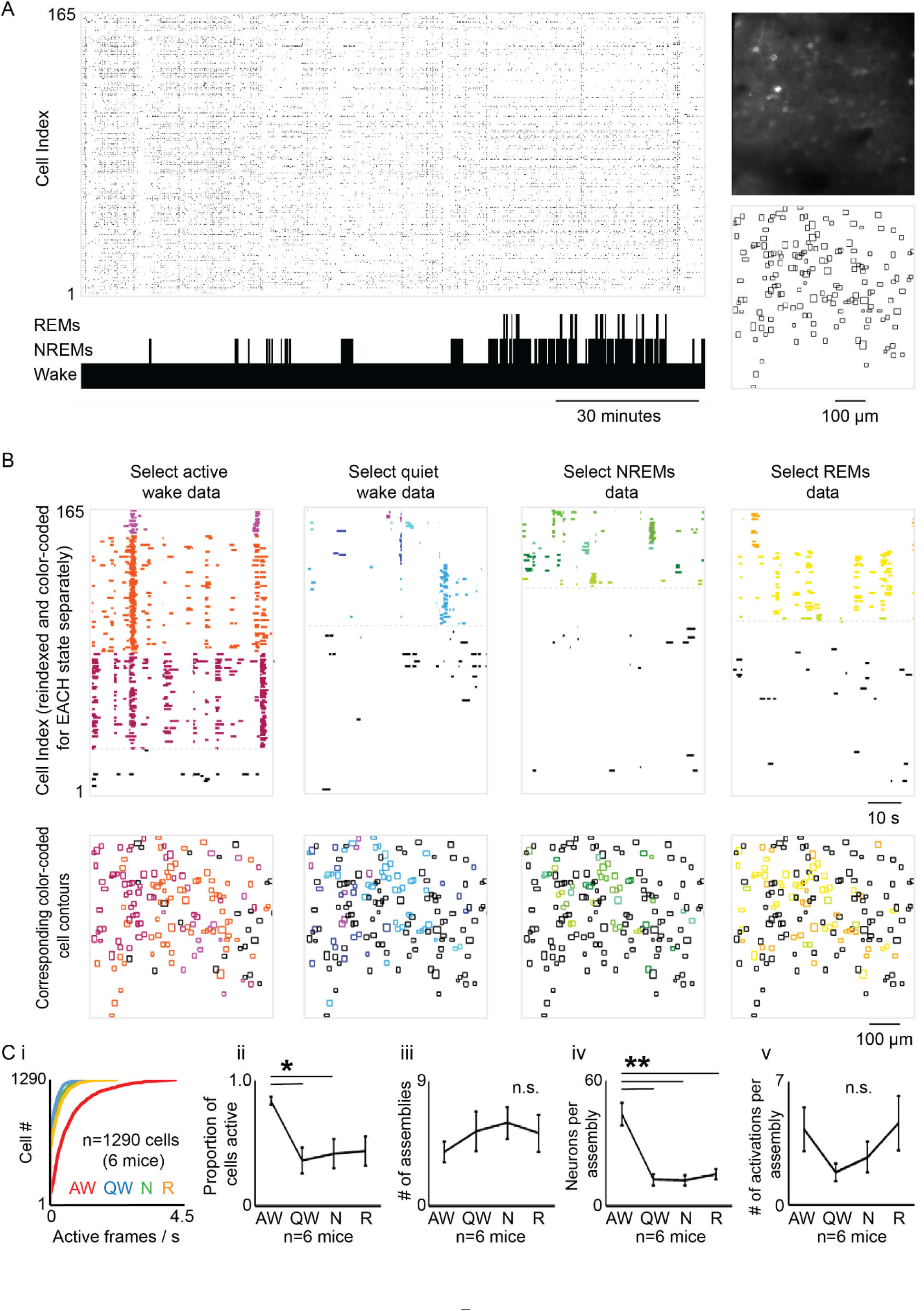
Detection of vigilance state-specific local microcircuit-level assembly activity in sensorimotor cortex. (**A**) Example of a full activity raster taken from an imaging experiment (black=active). The hypnogram at bottom indicates the behavioral state at a given time point. Right. An imaging frame taken from the experiment showing the field of view in the sensorimotor cortex is included at top. Neuron contours obtained from the field of view are shown at bottom. (**B**) Sample vigilance state-specific hierarchical clustering results obtained from select data originating from the raster shown in (**A**). For each vigilance state, significant assemblies that were identified have had their constituent neurons reindexed and color-coded for clarity. Cells plotted below the faded grey line were not clustered into a significant assembly. Bottom. Vigilance state-specific neuron contour plots with color-coding corresponding to the raster plots above. (**C i-v**) Group level analyses of basic cortical neuron and assembly activity from select data obtained for each vigilance state. (**i**) Plots of firing rate distributions for all neurons pooled across experiments. See Table S6 for statistical analysis of group level mean firing rate values derived from this data. (**ii**) Mean proportion of cells that were active for at least 1 frame during select data. (**iii**) Mean number of significant assemblies identified. (**iv**) Mean number of constituent cells per assembly. (**v**) Mean number of assembly activations. (n=6; n.s.=not significant, \**P*<0.05, \*\**P*<0.01, one-way repeated measures ANOVA with Tukey’s multiple comparisons post-hoc test (**ii-iv**) / Kruskal-Wallis test with Dunn’s multiple comparisons post-hoc test (**v**)). All group-level data are presented as mean ± SEM.

We next turned our focus to the analysis of vigilance state-specific local microcircuit-level cortical assembly activity. To evaluate the presence of significant assembly activity we utilized an agglomerative hierarchical clustering method that, while time-intensive, had the significant advantage of not requiring predefinition of the final cluster number (see supplementary methods). For each experiment, clustering was performed on raster data for each state individually and clusters were only considered as assemblies if they were composed of no fewer than 5 constituent neurons. Using this procedure, we were able to identify significant assembly activity in data from every vigilance state of each experiment (Fig. 2B). There was no difference between vigilance states in the number of assemblies identified (Fig. 2C, iii); however, there were on average significantly more neurons per assembly during AW relative to QW, N, and R periods, which were indifferent from one another (Fig. 2C, iv). We then measured assembly activity during the different vigilance states. An assembly was considered active during a given frame if at least 3 of its constituent neurons were simultaneously active; consecutive frames as well as frames that were not separated by a period where no constituent cells were active were not counted. This approach did not reveal a significant difference in the number of assembly activations between any of the vigilance states tested (Fig. 2C, v).

### Identification of global EEG microstates from multi-EEG recordings during different vigilance states

While our result of having found no clear consistent differences in local microcircuit-level cortical assembly structural characteristics between QW, N, and R behaviors could be inconsistent with the idea that they are critically involved in consciousness, we hypothesized that there might be a difference in the coordination of assembly activity with global brain dynamics. Indeed, recent evidence has indicated that, while local patterns of general network activity are preserved (Hudetz et al., 2016; Lewis et al., 2012), disrupted functional connectivity and coordination between distant cortical regions is a hallmark of the anesthetized brain (Lewis et al., 2012; Barttfeld et al., 2015; Hudetz et al., 2015). An established approach that has provided much insight into macroscale brain dynamics involves analyzing how the relationship between EEG recordings obtained simultaneously from widespread cortical locations changes over time (Lehmann et al., 1987; Pascual-Marqui et al., 1995; Mégevand et al., 2008; Michel and Koenig, 2018). More specifically, this procedure involves clustering topographical EEG potential profiles during moments of peak variability between the EEG signals, and has consistently demonstrated that global EEG activity dynamically shifts between a relatively small number of quasi-stable configurations. Termed ‘EEG microstates’, such dynamics on the global EEG scale have attracted significant interest in recent years due to their potential involvement in general cognition and conscious experience (Lehmann et al., 1998; Milz et al., 2016; Seitzman et al., 2017; Santarnecchi et al., 2017). We therefore next sought to investigate the coordination of local microcircuit-level cortical assembly activity with global brain dynamics in a subset of 3 mice that had been implanted with multi-EEGs (Fig. 3A). As a first step, we clustered topographical EEG potential profiles from data corresponding to periods of peak variation between EEGs during the same time periods used for imaging-derived assembly analysis in the sensorimotor cortex for QW, N, and R. AW periods were not analyzed due to the strong possibility of contamination of EEG signals by movement-related artifacts. Significant clustering of EEG topographies was found for all vigilance states tested in each experiment, with an optimal cluster number of 4 being consistently indicated for all vigilance states in each experiment (Fig. S1A-B). The ordered sets of EEG potential profile maps (microstate maps 1-4) derived from the optimal cluster definitions were highly similar between different vigilance states within an experiment (mean Pearson ***r*** values: QW-N = 0.79 ± 0.08; QW-R = 0.91 ± 0.01; N-R = 0.83 ± 0.07) (Fig. S2 A-B; Table S1). Microstate maps were also positively correlated between experiments (mean inter-experiment Pearson ***r*** values: QW = 0.80 ± 0.07; N = 0.68 ± 0.10; R = 0.41 ± 0.10) (Fig. S2 A-B; Table S2). To gain insight into the dynamics of EEG microstates, we next correlated each microstate map with instantaneous EEG potentials at each time point of the data segments (Fig. 3B). Microstates were found to cycle dynamically during all states; however, spectral analysis revealed that the primary frequencies at which this cycling occurred differed depending on vigilance state (Fig. 3C). Spectra for both QW and N were dominated by peaks in the delta (1-4 Hz) frequency range (mean dominant frequency: QW = 3.0 ± 0.4 Hz; N = 3.6 ± 0.3 Hz) although higher frequency dynamics associated with transient spindle oscillations (9-15 Hz) during select N data were sometimes observed (Fig. S3). In contrast, spectra for R data peaked in the theta (4-10 Hz) range (mean dominant frequency: R = 6.3 ± 0.2 Hz). Correlation analysis between microstate dynamics and each individual EEG channel indicated that no single EEG exerted disproportionate influence over microstate dynamics (Table S3). We also correlated microstate dynamics versus the average complex wavelet transform across all EEGs at each time point and found that microstate dynamics were not influenced by changes in global oscillation strength for any of the standard frequency bands tested (Table S4). Thus, our results are in general agreement with those described previously (Lehmann et al., 1987; Pascual-Marqui et al., 1995; Mégevand et al., 2008; Michel and Koenig, 2018; Lehmann et al., 1998).

**Fig. 3.**
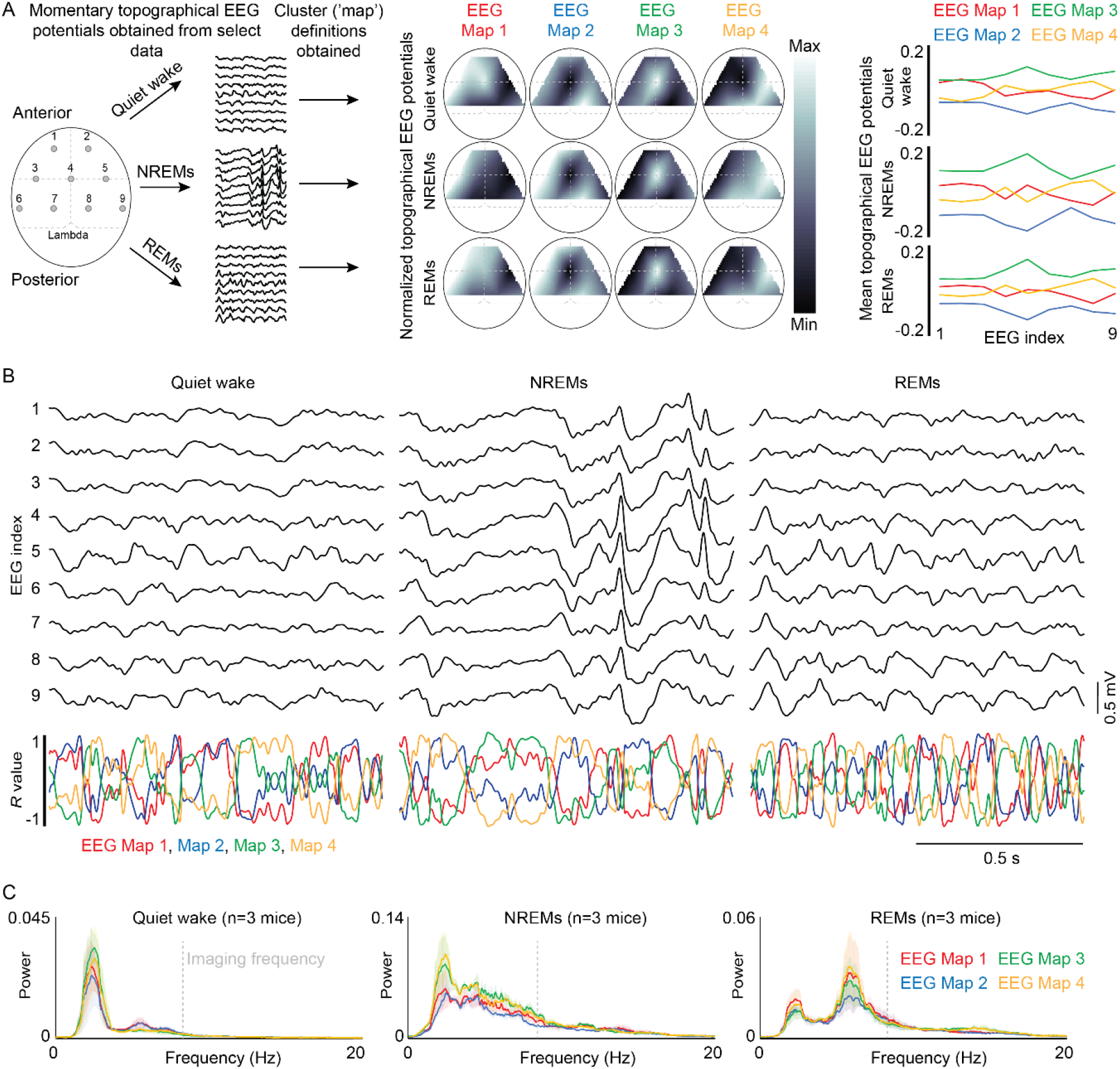
Identification of global EEG microstates from multi-EEG recordings during different vigilance states. (**A**) Schematic of procedure used for EEG microstate detection. Left. Multi-EEG signals originating from EEG electrodes widely spread over the cortical surface were obtained for the same select quiet wake, NREMs, and REMs time periods used for imaging data analysis. Note that the imaging window is not included in the dorsal-view schematic for clarity. The topographical EEG potential profiles were then clustered at time points corresponding to periods of peak variation between EEGs, resulting in the identification of significant clusters (see figure S4 and supplemental methods for details). Middle. Example normalized topographical EEG profile ‘microstate maps’ derived from optimal significant cluster definitions identified in prior steps for one experiment. Data are scaled according to the max-min difference for each map individually. Right. Linear plots of the same topographical EEG maps to provide direct comparison of values for each map. (**B**) Top. Expanded example of multi-EEG data plots for each vigilance state analyzed. Bottom. Corresponding plots of Pearson correlation ***r*** values calculated between momentary multi-EEG topographical profiles and microstate map templates (identified in (**A**)) used to show the relationship between multi-EEG data and EEG microstate dynamics. (**C**) Group level spectral analyses of identified global EEG microstate dynamics (Pearson correlation ***r*** values calculated between momentary multi-EEG topographical profiles and microstate map templates at each data point). The dashed grey line in each vigilance state-specific figure indicates the frequency of imaging data acquisition for reference.

### Microcircuit-level assembly activity in sensorimotor cortex is coordinated with global EEG microstates during quiet wakefulness

To investigate the potential relationship between EEG microstate dynamics and local microcircuit-level assembly activity in the sensorimotor cortex, we next calculated the locations of estimated cortical assembly activation start points during QW and N separately for each experiment (Fig. 4A-B). Start points were defined for each assembly activation as the time point at which the first constituent neuron became active. Values for R were not calculated because the dominant frequency of microstate dynamics during this vigilance state was not less than the Nyquist frequency of imaging data (half the imaging frame rate (8 Hz)), which would have been susceptible to under sampling as a result. The correlation coefficient ***r*** values calculated for each EEG microstate map corresponding to the 125 ms (~ imaging frame rate) period before and 125 ms period after the estimated start point were then determined for each assembly activation, and average values were calculated for each vigilance state tested. We found that ***r*** values corresponding to EEG microstate maps 1 and 2, associated with relatively increased potentials in frontal and posterior EEGs, in the 125 ms period following estimated assembly activations were significantly higher relative to the preceding 125 ms time window during QW. In contrast, no such differences were found for assembly activations occurring during N periods (Fig. 4B-C; Fig. S4, top-middle). To ensure that this result was not due to under sampling of imaging data during transient spindle-associated periods of increased frequency EEG microstate dynamics during N periods, we next automatically detected spindle activity in all EEGs (Fig. S3). Spindle activity coincidence, defined as significant spindle activity being detected in at least 1 EEG channel during the −125: +125 ms peri-assembly activation period, was only found to occur in a minority (11 %) of assembly activations; the removal of these data points did not significantly alter our results (Fig. S4, bottom).

**Fig. 4.**
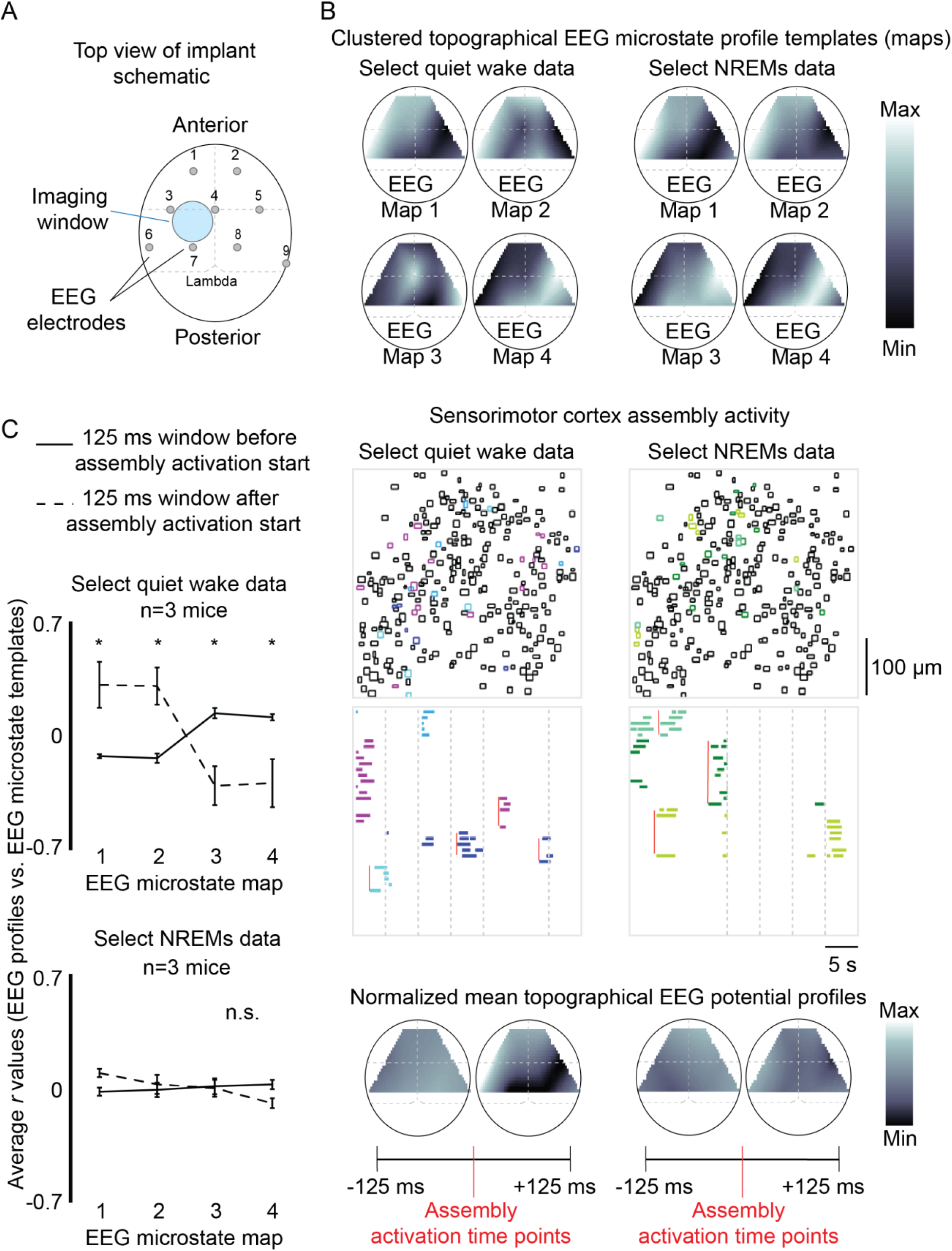
Microcircuit-level assembly activity in sensorimotor cortex is coordinated with global EEG microstates during quiet wakefulness. (**A**) Dorsal view schematic of the implant which allows for simultaneous imaging of the sensorimotor cortex and multi-EEG recording. (**B**) Example showing vigilance state-specific correspondence between EEG microstate dynamics and microcircuit-level assembly activity in the sensorimotor cortex. Top. Sample EEG microstate template maps generated for select quiet wake and NREMs data. Data are scaled according to the max-min difference for each map individually. Middle. Color-coded contour maps and corresponding spike rasters obtained from the same select data periods as EEG microstate maps shown at top. To assess the potential relationship between microcircuit-level assembly activity and global EEG microstate dynamics during quiet wakefulness and NREMs, the estimated start points of cluster activations were obtained (indicated by red vertical lines in the raster plots, see text for details). Assembly activations with start points located within 125 ms of a temporal break in the data (indicated by vertical dashed grey lines) were not used for analysis. Bottom. The mean topographical EEG potentials were then calculated for the 125 ms windows immediately prior to and after each valid assembly activation. The averaged topographical potential values are shown in the plots, which have been normalized with max/min limits according to the data set with the greatest variation within its own map (select quiet wake +125 ms). Monotone plots during the 125 ms windows both preceding and following valid assembly activations in NREMs data are indicative of relatively reduced correspondence between microcircuit-level assembly activation time points and EEG microstate dynamics as compared to quiet wake. (**C**) Group level analysis of Pearson correlation ***r*** values calculated between momentary multi-EEG topographical profiles and microstate map templates at each data point within the +/- 125 ms windows surrounding assembly activation start points. (n.s.=not significant, \**P*<0.05, two-way repeated measures ANOVA with Holm-Šídák’s multiple comparisons post-hoc test). All group-level data are presented as mean ± SEM.

## Discussion

Although prior work has suggested that local microcircuit-level cortical assemblies may be fundamental units of consciousness, there has been a lack of detailed information about their structural characteristics across the full spectrum of natural vigilance states. Here, we performed a comparative characterization of local microcircuit-level assemblies in the sensorimotor cortex of mice during periods of AW, QW, N, and R. We report that there was no significant difference in either the mean number of significant assemblies identified or in the mean assembly activity rates between any of the vigilance states tested. Furthermore, while assemblies occurring during AW were on average composed of more cells, there were no such differences found between assemblies occurring during QW, N, or R, despite the latter two vigilance states being associated with significantly reduced levels of consciousness relative to QW. However, when the potential coordination between assembly activity and global brain dynamics (assessed through EEG microstate analysis) was tested, we found that assembly activations were associated with an increase in the occurrence of EEG microstates corresponding to relatively high potentials in frontal and posterior EEG electrodes during periods of QW, but not N (AW and R were not tested for logistical reasons). These results suggest that the coordination of local microcircuit-level cortical assembly activity with global brain dynamics could be key for sustaining a stable state of consciousness, whereas the mere presence of cortical assemblies is necessary (Wenzel et al., 2019), but not sufficient. These results are also in line with prior experiments which have demonstrated that, while local patterns of general network activity can be preserved (Hudetz et al., 2016; Lewis et al., 2012), disrupted coordination between distant cortical regions is observed during anesthesia-induced unconscious states (Lewis et al., 2012; Barttfeld et al., 2015; Hudetz et al., 2015; but see Wenzel et al., 2019).

In this study, we employed EEG microstate analysis as a method to gain insight into global brain dynamics. While this analysis is derived from cortical EEG signals, the brain-wide relevance of EEG microstate dynamics has been suggested by prior studies that associated them to resting state networks in dual EEG-fMRI experiments (Musso et al., 2010; Britz et al., 2010; Britz et al., 2010; Yuan et al., 2012), as well as studies which used source localization techniques to reveal multiple underlying contributors to the EEG signal (Custo et al., 2017; Milz et al., 2017; Pascual-Marqui et al., 2014). Furthermore, a similar ‘EEG microstate’ analysis approach was applied to data obtained from high-density electrode arrays implanted in the prefrontal cortex, striatum, and ventral tegmental area in a recent study utilizing a rat model (Mishra et al., 2021). The authors reported that each brain region produced dynamics similar to those of more traditional EEG studies, and that these dynamics were synchronized to some extent across the different regions. Cumulatively, these studies suggest that EEG microstate dynamics may be a direct reflection of transient alterations in the functional connectivity between a variety of different brain regions, although currently little is known of their direct relation to underlying neural activity (Michel and Koenig, 2018). Thus, our study also provides important novel insight by demonstrating the relationship between local microcircuit-level cortical assembly activity and EEG microstate dynamics during periods of restful wake. More generally, our data suggests that EEG microstates could be a mechanism by which local microcircuits are able to coordinate across relatively distant brain regions in the awake brain. Our present work thus highlights a new promising direction of research with potentially significant implications considering, aside from their potential role in general cognition and conscious experience (Lehmann et al., 1998; Milz et al., 2016; Seitzman et al., 2017; Santarnecchi et al., 2017), the possible association between aberrant microstate dynamics and neuropsychiatric disorders, most notably schizophrenia (Irisawa et al., 2006; Kikuchi et al., 2007; Koenig et al., 1999; Lehmann et al., 2005; Nishida et al., 2013; Strelets et al., 2003; Andreou et al., 2014; Tomescu et al., 2014; Kindler et al., 2011) among others (Nishida et al., 2013, Grieder et al., 2016; Drissi et al., 2016; Kikuchi et al., 2011; Gschwind et al., 2016; Corradini et al., 2014; Gao et al., 2017; Zappasodi et al., 2017).

In conclusion, our results suggest that the coordination of cortical assembly activity with global brain dynamics could be a key factor of sustained conscious experience. More generally, this data provides novel evidence that EEG microstates could play an important mechanistic role in promoting coordination and temporary binding between local activity patterns in physically segregated neural populations at the microcircuit-level in the awake resting brain.

## Acknowledgements

RB was supported by The International Human Frontier Science Program Organisation (HFSP) postdoctoral fellowship #LT000835/2018-L while completing this research. This project also received support from ERC Neuropioneers (Grant Number 646925 - ERC-2014-CoG), the Fondation Bettencourt, and ERC Hope (Grant NUMBER 95133 - ERC Synergy Grants 2020).

## Author contributions

RB conceived and designed the research project, completed surgical implantation procedures and subsequent experiments in adult mice, performed all data analysis, and wrote the manuscript. RD performed intraventricular virus injections in mouse pups. RC established the laboratory in which experiments were completed, supervised and provided guidance for the study, and provided financial support.

## Method details

### Experimental model and subject details

All experiments were performed in mice under the guidelines of the French National Ethics Committee for Sciences and Health report on “Ethical Principles for Animal Experimentation” in agreement with the European Community Directive 86/609/EEC (Apafis#28.506). When not being used in experiments, mice were housed in cages on a 12h:12h light:dark light schedule (lights on at 20:00 h) and had access to food and water *ad libitum*. Mice were housed in individual cages following weaning at 3 weeks age.

#### Virally-mediated expression of calcium indicators

*In-vivo* calcium imaging experiments were facilitated through cortical expression of GCaMP6s via the viral vector AAV1-hSyn-GCaMP6s.WPRE.SV40 (a gift from Douglas Kim & GENIE Project (Addgene viral prep #100843-AAV1; http://n2t.net/addgene:100843; RRID:Addgene_100843)). To achieve stable widespread GCaMP6s expression long-term throughout the cortex, we used an intracerebroventricular injection protocol adapted from previously published methods (^13-14^). On post-natal day 0, Swiss mouse pups were anesthetized on ice for 3 – 4 min and 2 μL of viral solution (titration at least 1 x 1013 vg / mL) were injected in the left lateral ventricle whose coordinates were estimated at 2/5 of the imaginary line between the lambda and the eye at a depth of 0.4 mm.

#### Implantation of electrodes and imaging window

At approximately 3 months age, transduced male mice were anesthetized with isoflurane (5 % induction, 1-2 % maintenance) and placed into a stereotaxic frame. Skin covering the top of the skull was removed and the skull surface was cleared of all connective tissue. The head position was subsequently adjusted so that the bregma and lambda were located in the same horizontal plane. The top of the skull was then lightly scored with a drill and a thin layer of dental cement (C&B Metabond, Parkell, Edgewood, NY) was applied to the skull surface. In all of the mice, a ~ 250 μm diameter hole was drilled through the skull (anterior-posterior (AP) −2.67, medial-lateral (ML) +1.54, all values in mm relative to bregma) and a stainless-steel EEG screw (Antrin Miniature Specialities, Inc.) with an insulated tungsten wire lead (A-M systems; item # 795500) soldered to the screw head was inserted just until secure to avoid damaging the underlying dura. 2 additional screws were similarly placed in the bone above the frontal cortex and cerebellum to serve as ground and reference electrodes, respectively. In order to investigate the dynamics of global brain EEG microstates, 8 additional EEG screws (multi-EEGs) were added at the following coordinates in a subset of the mice: (AP +2.67, ML ± 1.54; AP 0.00, ML 0.00, ± 3.08; AP −2.67, ML −1.54, ± 4.62). In addition, an insulated stranded wire (Medwire) was inserted into the nuchal muscle to serve as an electromyogram for recording of postural muscle tone. To facilitate 2-photon calcium imaging of cortical neural activity, a 3 mm diameter circular portion of the skull centered over the sensorimotor cortex of the left hemisphere (center coordinates: AP −0.75, ML −1.75) was then carefully removed. Next, a small amount of clear silicon polymer (Kwik-sil (World Precision Instruments)) was applied to the top of the dura and a 3 mm diameter glass window was subsequently placed on top. A metal bar, required for head-fixation of mice during later experiments, was placed at the back of the skull behind the cerebellar reference screw. Mill Max pins (Duratool corporation) soldered to the wire leads originating from the EEG, EMG, ground, and reference electrodes were then fixed in a precise configuration on top of the metal bar; pins were oriented towards the rear of the mouse at a vertical angle of 45° to keep enough space for the objective and also prevent the headstage preamplifier from obstructing the mouse when head fixed. As a final step, all electrodes were insulated and the entire implant was secured with additional dental cement. Mice were then returned to their home cage and given buprenorphine (0.1 mg/kg) to help with pain management during the initial recovery period. Mice were given 3 weeks to rest in order to ensure a full recovery from the surgery. At this time, the quality of the implanted electrodes and imaging field of view (FOV) was assessed; of 13 total mice that underwent the procedure outlined above, 6 were excluded from further experimentation at this point due to a poor FOV signal to noise ratio (due to low levels of GCaMP6s expression and / or the presence of debris or air pockets under the imaging window which interfered with the fluorescent signal below). In the remaining 7 mice, 4 of which had multi-EEG implants, the behavioral habituation procedure was started.

#### Habituation to head fixation

Once fully recovered from the electrode and imaging window implant procedure, habituation sessions for prolonged head fixation commenced. The start of each session occurred ~ 1 h before the start of the light cycle to encourage sleeping behavior. Using the metal bar attached to the head of the mouse during the prior surgery, mice were head fixed underneath the 2-photon microscope on a non-motorized treadmill that allowed them to run at will. Light in the room was kept to a minimum to prevent contamination of the imaging signal. For each mouse, a single dedicated tread was used for the entire duration of a given habituation and recording procedure. The head position was customized for each mouse to ensure that they would be able to sleep in a sphinxlike position on the tread while leaving enough space to allow for running. Mice participated in 1 training session per day, with the duration of head fixation being gradually increased across 12 successive sessions to a maximum duration of 4.5 h: 5 min, 15 min, 30 min, 45 min, 1 h, 1.5 h, 2 h, 2.5 h, 3 h, 3.5 h, 4 h, 4.5 h. Once the maximum duration of 4.5 h was reached, the daily head fixation time remained constant until the experiment end point was reached. This procedure was tolerated well by all mice used in this study; all but 1 of the 7 remaining mice demonstrated a consistent full sleep-wake cycle (wakefulness, NREMs, REMs) while head fixed that was relatively stable from day-to-day ~4 weeks after starting habituation. The remaining mouse tolerated the procedure well; however, as there was little sustained (> 5 s continuous) sleep activity, it was excluded from any further data acquisition and analysis. Sessions were completed daily throughout the duration of an experiment for a given mouse (habituation and subsequent data acquisition), unless mice began to demonstrate signs of fatigue (significant (>50 %) decrease in sleep quantity and weight loss (more than 5 %) relative to prior day), in which case they were allowed to rest at least 1 day or until weight recovered (typically 1-2 days), respectively. ~2 weeks of data acquisition was typically required for a given mouse to obtain a single experiment with optimal quality imaging and EEG data as a result of common logistical issues related to imaging (primarily loss of imaging FOV mid-experiment due to bubble formation or a shift in the FOV position that could not be corrected post-hoc). For each mouse, only data from a single optimal experiment was used for subsequent data preprocessing and analysis steps. Finally, to assess the potential impact of head fixation on sleep quality, the day following completion of data acquisition, the treadmill apparatus was removed from the 2-photon imaging platform and replaced by an empty cage with woodchip bedding. The mouse was placed un-head fixed in the cage with the head stage pre-amplifier tether attached, and was subsequently recorded under otherwise identical experimental conditions (i.e., at the same time, for the same duration, under the same lighting conditions, and with the microscope scanner active). Once this endpoint was reached, mice were utilized for experiments in an unrelated project.

#### Electrophysiological recording

Electrophysiological recordings were done during all phases of the experiment (head-fixation habituation in order to track habituation progression as well as during imaging data acquisition experiments in fully habituated mice). For each recording, a head stage pre-amplifier (Neuralynx, Boseman, Montana, USA) tether was attached to the Mill-max connector pins at the rear of the head of the mouse immediately after being head fixed into the 2-photon imaging setup. Data from all electrodes were subsequently amplified by the head stage pre-amplifier before being digitized at 16000 Hz using a digital acquisition system (Neuralynx, Boseman, Montana, USA) and saved to a hard disk. Tread movement was also captured through a video recording synchronized to the electrophysiology data.

#### 2-photon imaging of cortical neural activity

To keep the size of data sets manageable and minimize the potential for photobleaching, imaging data acquisition was limited to the 2nd half of the experiment (~ 2.25 h in duration, hour 2.25-4.5 of experiment) when sleep activity was generally highest. At the start of this period, a 16 x immersion objective (NIKON, NA 0.8) connected to a GaSP PMT (H7422-40, Hamamatsu) was positioned over the imaging region of interest (field of view (FOV)) within the sensorimotor cortex at an imaging depth corresponding to ~150-250 μm below the pial surface. A series of 25 movies (2500 frames per movie acquired at 8 Hz, FOV size = 400 x 400 μm acquired at a resolution of 200 x 200 pixels) was obtained using a single beam multiphoton-pulsed laser scanning system coupled to a microscope (TriM Scope II, LaVision Biotech) and Ti: sapphire excitation laser (Chamelon Ultra II, Coherent) operated at 920 nm. GCamp6s fluorescence was isolated using a bandpass filter of 510 nm/25nm. Images were acquired from the PMT signal using Imspector software (LaVision, Biotech) and subsequently saved to a hard disk. There was no evidence of photo-toxicity or significant bleaching as a result of the above imaging procedure. To synchronize imaging data with electrophysiological data, a TTL timestamp signal was sent from the imaging acquisition software to the digital electrophysiology acquisition system at the onset of each imaging frame scanning cycle.

#### Data analysis

All following data analysis was performed using custom written scripts in Matlab (The MathWorks, Inc.).

#### Vigilance state architecture analysis

For quantitative and qualitative analysis of sleep and wake activity of each experiment, raw EEG and electromyogram data files were first imported into Matlab and downsampled to 1000 Hz. Data was then manually plotted and the major phases of the sleep-wake cycle occurring throughout the entire recording session were scored in 5 s epochs using the fast Fourier transform (FFT; ‘fft’ function in Matlab) of the signal recorded from the right parietal EEG (AP −2.67, ML +1.54) in addition to the nuchal EMG signal (Boyce et al., 2016). The procedure was as follows: epochs of wakefulness were identified by a binned ‘theta’ (4-10 Hz, ‘θ’) to ‘delta’ (1-4 Hz, ‘δ’) power ratio greater than 1 and bursts of relatively high-amplitude movement-associated EMG activity (typically > 0.1 mV and > 1 s in duration). Epochs of NREMs were identified by a binned θ / δ power ratio < 1 and a lack of high-amplitude EMG activity. Transient spindle (9-15 Hz) oscillations were also observed periodically during NREMs. Epochs of REMs followed and preceded NREMs and wakefulness, respectively, and were identified by a binned θ / δ power ratio greater than 1 and a completely flat EMG signal except for relatively brief and phasic (< 1 s in duration) periods of high-amplitude EMG activity associated with muscle twitches. While many state transitions were relatively gradual, occurring over the course of several epochs (e.g., from wakefulness to NREMs, NREMs to REMs) and thus allowing the above criteria to be applied with ease, epochs during which clear and relatively abrupt state transitions occurred (e.g., REMs to wakefulness) were scored as being the state that occupied the majority of the epoch. Once scoring was completed, the data was manually reviewed and crosschecked with video recordings to ensure a perfect correspondence between overt mouse behavior (e.g., movement, grooming) and the scored vigilance state data (hypnogram). Total state proportions and average state durations for each experiment were calculated based on the hypnograms generated using the above procedure.

For EEG microstate and imaging data analysis, periods of wakefulness were further segregated into ‘active wake’ or ‘quiet wake’ according to the presence or absence of activity bursts in the EMG signal, respectively. To ensure that periods of early transitional sleep were not accidentally identified as quiet wake, only periods of quiet wake completely free of any events reminiscent of large slow wave and spindle-like activity that were flanked by overt periods of active wakefulness were selected.

#### Spectral analysis of EEG data

For analyses that prioritized frequency resolution (creation of basic EEG spectrograms, spectral analysis of EEG microstate dynamics), vigilance state-specific spectral analysis of EEG data was completed in Matlab using the mtspecgramc function from the Chronux signal processing toolbox (parameters: window size = 5 s, step size = 5 s, tapers [3 5]) (Bokil et al., 2010).

#### Wavelet analysis of EEG data

For analyses that prioritized temporal resolution (correlation of specified frequency bands with EEG microstate dynamics), vigilance state-specific Morlet wavelet analysis of EEG data was completed in Matlab using the ‘cwt’ function. The absolute values of the resulting complex wavelet transforms were then calculated for each time point, and data were binned according to the following definitions: ‘delta’ (1-4 Hz, δ); ‘theta’ (4-10 Hz, θ); ‘alpha’ (9-15 Hz, α); ‘beta’ (15-30 Hz, β); ‘gamma’ (30-50 Hz). Values originating from datapoints within 0.5 s of a temporal break in the data were ignored.

#### EEG microstate analysis

For identification of EEG microstates, we used a protocol similar to a well-established procedure that has been described in detail previously (Pascual-Marqui et al., 1995; Michel and Koenig, 2008). EEG traces from each experiment were first bandpass filtered from 1-50 Hz using the ‘filtfilt’ function in Matlab. The ‘global standard deviation’ (GSD), the standard deviation of all EEG signals, was then calculated for each data timepoint and the indices of peak locations in the GSD were subsequently found using the ‘findpeaks’ function in Matlab. For each experiment, equal amounts of quiet wake, NREMs, and REMs data that were free of activity bursts in the EMG were then identified. Periods of active wake were not analyzed due to the high potential for contamination of EEG signals with movement artifacts. The amount of data available for each state was therefore limited by the state with the least amount of data available (typically REMs). Valid data points for other states that had to be excluded as a result were removed at random. The EEG profile maps, the instantaneous potential of each EEG signal, associated with the indices of each identified peak in the GSD occurring during identified EMG noise-free quiet wake, NREMs, and REMs periods were then clustered separately as well as in combination (i.e., select quiet wake, NREMs, and REMs periods combined) using the k means clustering ‘kmeans’ function in Matlab (parameters: maximum number of clusters=10, number of iterations per clustering step=100), followed by silhouette analysis using the Matlab ‘silhouette’ function. For statistical evaluation of clustering results, the above procedure was repeated 100 x using shuffled surrogate data which was generated by shifting each EEG channel individually relative to their initial position by a random amount. The resulting outcomes of clustering were highly similar both for different states (including all states combined condition) within a single experiment as well as between different experiments, consistent with prior reports (Pascual-Marqui et al., 1995; Michel and Koenig, 2008). Therefore, to best facilitate comparison of microstate dynamics between different states, the optimal cluster number that was subsequently used for state-specific microstate analysis was set at that which produced an explained variance closest to 90 % in clustering of combined (select quiet wake, NREMs, and REMs) data. This approach was used instead of taking the cluster number associated with the peak silhouette value in order to capture the maximum reasonable amount of variation in the data. For each individual state (select wake, NREMs, and REMs) of each experiment, the clustering results associated with the optimal cluster number determined using the aforementioned procedure were statistically significant (silhouette value higher value than that from at least 95/100 of the shuffled surrogates). As a final step, EEG profile template ‘maps’ were generated for every individual cluster by averaging, for each EEG, the potentials associated with all constituent timepoints of the cluster. Potential maps corresponding to each cluster were then reindexed according to their general features which were consistently observed both within (i.e., different vigilance states within the same animal) and between experiments (i.e., different mice). Potential ‘Map 1’ was characterized by relatively increased potentials in the frontal and posterior electrodes. ‘Map 2’ was also characterized by relatively increased potentials in the frontal and posterior electrodes; however, the profile was shifted more negatively relative to those of ‘Map 1’. ‘Map 3’ was characterized by relatively increased potentials in the middle EEG electrodes, while ‘Map 4’ was also characterized by relatively increased potentials in the middle EEG electrodes but shifted more negatively relative to those of ‘Map 3’.

#### Analysis of spindle activity in multi-EEG data

Spindle activity in EEG recordings was completed using a method similar to one used previously (Boyce et al., 2016). Analysis was completed on temporally continuous periods of select NREMs data (the same periods used for EEG microstate analysis during NREMs) using non-overlapping 100 ms windows; the following procedure was completed for each EEG channel individually. For each window, EEG data was first bandpass filtered (9-15 Hz) using the ‘filtfilt’ function in Matlab. The data was then rectified and the sum of the absolute amplitude values was calculated. The mean and standard deviation (stdev) of absolute amplitude data values derived from all analysis windows was subsequently obtained and used to determine the threshold for spindle detection (mean + 3 stdev). All time points associated with values exceeding this threshold were considered to be locations of significant spindle activity.

#### Motion correction of imaging data

Raw imaging movies for each experiment were first loaded into Matlab, concatenated into a single 3-dimensional matrix, and subjected to motion correction using the NoRMCorre algorithm (Pnevmatikakis and Giovannucci, 2017). The procedure consisted of 2 rounds of rigid motion correction with successively stricter corrective parameters (bin size of 10 and maximum allowable shift of 30 for the first round; bin size of 2 with a maximum allowable shift of 15 for the second round). This was followed by 2 rounds of non-rigid motion correction, also with successively stricter corrective parameters (grid size of ¼ the imaging field of view, bin size of 3, and maximum allowable shift of 15 for the first round; grid size of 1/8 the imaging field of view, bin size of 2, and a maximum allowable shift of 10 for the second round). Following the above motion correction procedure, the final corrected movie was visualized with the un-corrected data and manually scanned to ensure the absence of significant overcorrections and / or other artifacts resulting from the algorithm.

#### Determination of neurons (regions of interest) in imaging field of view

The identification of regions of interest (ROIs) corresponding to neurons in the field of view (FOV) of motion corrected movies was completed using a semi-automated approach. Each motion corrected movie was manually played back frame-by-frame; during the playback, regions of interest (ROIs, i.e., areas of the field of view with calcium signals characteristic of healthy cortical neurons) were manually identified. When a ROI was identified, a pixel in the center of the ROI was manually selected as a ‘seed’. The calcium signal of this seed pixel throughout the course of the full movie was then z-scored, and the similarity of this data vector was compared to that for each pixel in the immediate vicinity (pixels within 10 μm of the seed pixel) via linear correlation using the ‘corr’ function in Matlab. Pixels whose activity during the movie was highly correlated (P<0.01) with that of the seed pixel were automatically included as part of the ROI. If necessary, the pixels assigned to the ROI were then manually adjusted; adjustment was largely limited to the removal of pixels clearly associated with processes, as well as those overlapping / in direct juxtaposition with other cells. The above process was repeated until all clearly identifiable ROIs had been registered. At this point, a trace was derived for each ROI by averaging the calcium signal across all constituent pixels for each movie frame.

#### Identification of transient activity in calcium traces

Following extraction of calcium traces from identified ROIs, identification of calcium transients from this data was next completed for each experiment. For this, each raw calcium trace obtained from imaging data was first smoothed with a Gaussian filter using the ‘smoothdata’ function in Matlab and a smoothing window size corresponding to 1 s of data (8 frames). The ‘findpeaks’ Matlab function was then used to identify the location (timepoint), peak prominence, and width of peaks occurring in the smoothed trace. The onset timepoints for each peak were then identified by finding the closest preceding upwards deflection in the differentiated smoothed trace, calculated using the ‘diff’ function in Matlab. Peaks corresponding to significant calcium transients were then automatically detected by determining those which had prominences greater than the set threshold value of 250 units. This threshold value was determined for the current experiments as follows; first, the mean and standard deviation of the raw calcium signal for all ROIs was calculated during quiescent periods of the recording (periods where the animal was not moving and no cells were active) for each mouse. The value of 250 units was the nearest whole value (50 unit resolution required to best facilitate manual verification of detected peaks described below) that was greater than 3 standard deviations above the average standard deviation calculated for the noisiest experiment. The raw and corresponding smoothed traces, along with the calculated locations of significant calcium transients, were then verified by plotting the data in an interactive figure and scanning through the entire trace. Manual corrections were able to be made at this time; these were largely limited to the exclusion of transients detected during periods contaminated with uncorrectable movement artifacts, as well as the inclusion of clear transients occurring in quick succession. Once detection and manual verification of transients was complete, the data was binarized into spike rasters; for each ROI, periods without significant transient activity were indicated with a 0, while the periods corresponding to significant transient activity (frames occurring between the onset and peak location calculated for each significant transient) were indicated with a 1.

#### Hierarchical clustering of cortical imaging data

To evaluate the presence of assemblies in cortical imaging data, we used a hierarchical clustering procedure similar to one previously described (Feldt-Muldoon et al., 2013). Despite the disadvantage of being a relatively time-consuming technique compared to other clustering methods that could be used on this data set, we opted to employ this clustering procedure on our imaging data due to the significant advantage of there being no need to predefine the number of clusters present. To enable a comparison of assembly activity between different vigilance states, equal amounts of active wake, quiet wake, NREMs, and REMs data were identified (note that for mice with multi-EEGs imaging data was synchronized with the same quiet wake, NREMs, and REMs data time points used for EEG microstate analysis described in the previous section). The amount of data available for each state was therefore limited by the state with the least amount of data available (typically REMs). For select imaging data obtained for each vigilance state within an individual experiment, the hierarchical clustering procedure was as follows. First, the 0-lag associated cross-correlation value for each ROI pair was calculated using the ‘xcorr’ function in Matlab and stored in a correlation matrix. The corresponding P value of the correlation between each ROI pair was then found through the use of a block-shuffling technique. This procedure, which was repeated 100 x for each ROI pair, involved fragmenting the spike rasters of both ROIs into ~ 5 s blocks and independently reassembling them in random order before calculating and storing the correlation value. Once this process had been repeated 100 x, a cumulative distribution function was created using the values obtained for each shuffle with the ‘makedist’ function in Matlab, and subsequently used to determine the P value of the correlation value found between the intact spike rasters of the ROI pair using the Matlab ‘cdf’ function. Once the initial correlation and corresponding P-value matrices had been created, the main clustering procedure began. The index of the most statistically significant positive correlation in the entire matrix was found and the two associated ROIs were ‘clustered’ together with the corresponding spike rasters being merged. The values in the correlation matrix associated with the individual ROIs were nullified, and correlation and P-values were calculated between the newly merged spike raster and all of the remaining spike rasters. The index of the most statistically significant positive correlation in the entire matrix was again found and the associated ROIs (including correlations involving rasters that were the product of the merging of data from multiple ROIs) were clustered together. This process was repeated until there were no remaining significantly correlated spike raster pairs. At this point, the indices of ROIs (cortical neurons) that had been clustered together were stored. Only clusters consisting of at least 5 ROIs were considered valid in order to minimize the potential for chance cluster activations to introduce noise in subsequent analysis.

### Quantification and statistical analysis

Unless stated otherwise within the text, statistical analysis was performed using GraphPad Prism software (Dotmatics). *P*<0.05 was considered statistically significant, and all group-level data are presented as mean ± SEM. No subjects which underwent experimental recordings, nor data values produced from resulting analysis, were excluded from this study. Where appropriate, the decision to use parametric vs non-parametric statistical measures was strictly dependent on the results of normality and equal means testing using the Kolmogorov-Smirnov and Bartlett’s tests, respectively. Specific statistical details can be found in the figure legends, as well as in the ‘Method details’ section for shuffling-based statistical analysis used for evaluation of clustering results.

## Supplemental information

**Fig S1:**
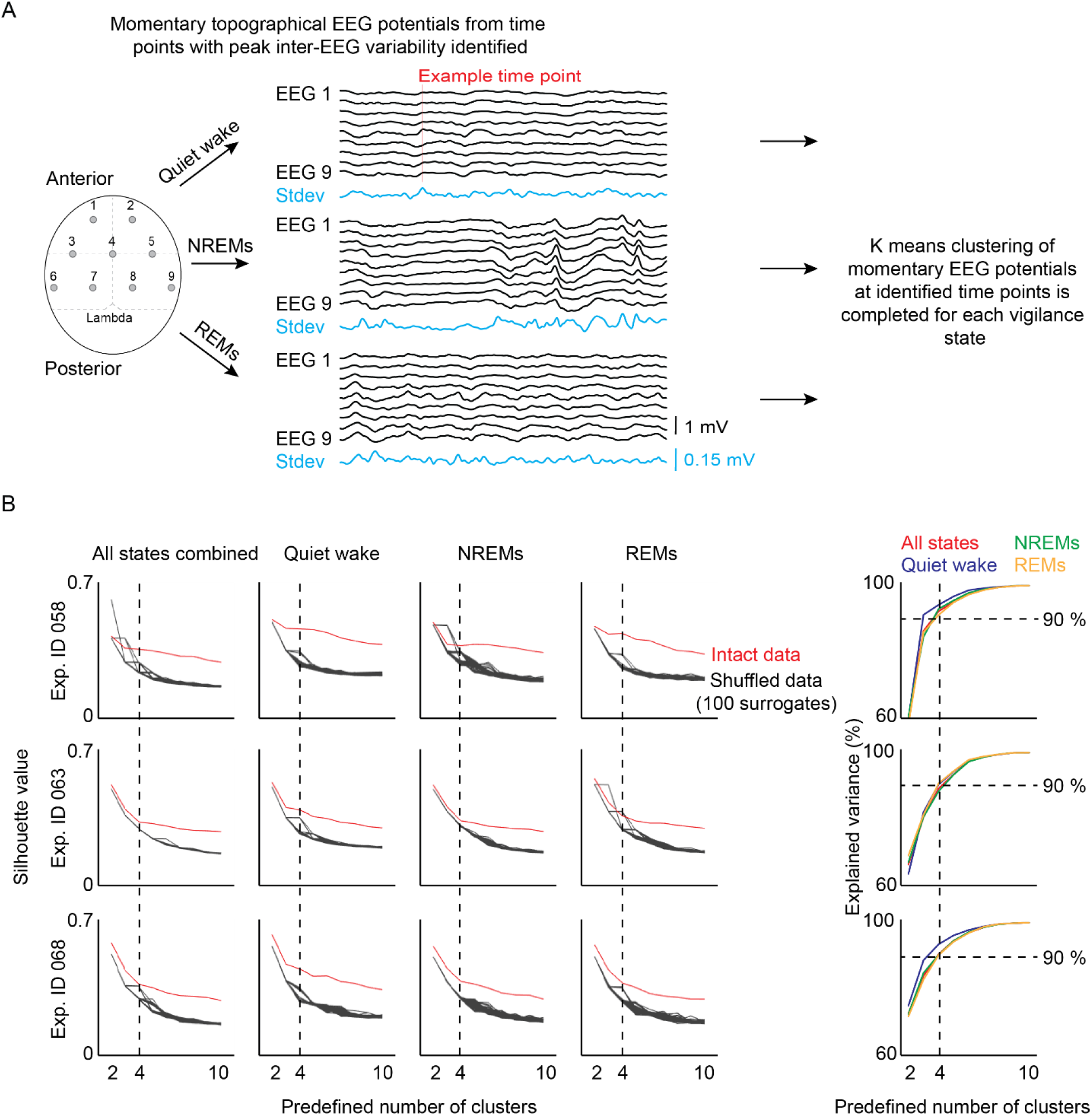
Identification of vigilance state-specific EEG microstates through k means clustering analysis of multi-EEG data. (**A**) Schematic of procedure used for EEG microstate detection. Multi-EEG signals were obtained for the same select quiet wake, NREMs, and REMs time periods used for imaging data analysis. Note that the imaging window is not included in the dorsal-view schematic for clarity. Time points corresponding to peaks in the variability (standard deviation) between EEGs were then identified for each vigilance state of a given experiment separately. Standard deviation data is plotted in blue below the sample traces. An example peak variation time point is indicated by the vertical red line in the quiet wake traces. Momentary multi-EEG potential profiles corresponding to all identified peak variation time points then underwent k means clustering to evaluate the presence of significant global EEG microstates. (**B**) Results of k means clustering of multi-EEG potential profiles are shown in scree plots for each vigilance state analyzed from each multi-EEG experiment (n=3 mice; experiment ID # 058, 063, 068). The red line indicates silhouette values obtained from intact data, while black lines indicate those obtained from 100 shuffled surrogates. Color-coded plots at right show the explained variance for each predefined cluster number for the ‘all states combined’ (QW + N + R) select data analysis condition. Clustering definitions generated using a cluster number that resulted in explained variance closest to 90 % were considered optimal for subsequent EEG microstate analysis (4 clusters in all experiments, for all vigilance states). See supplementary methods for more details.

**Fig. S2.**
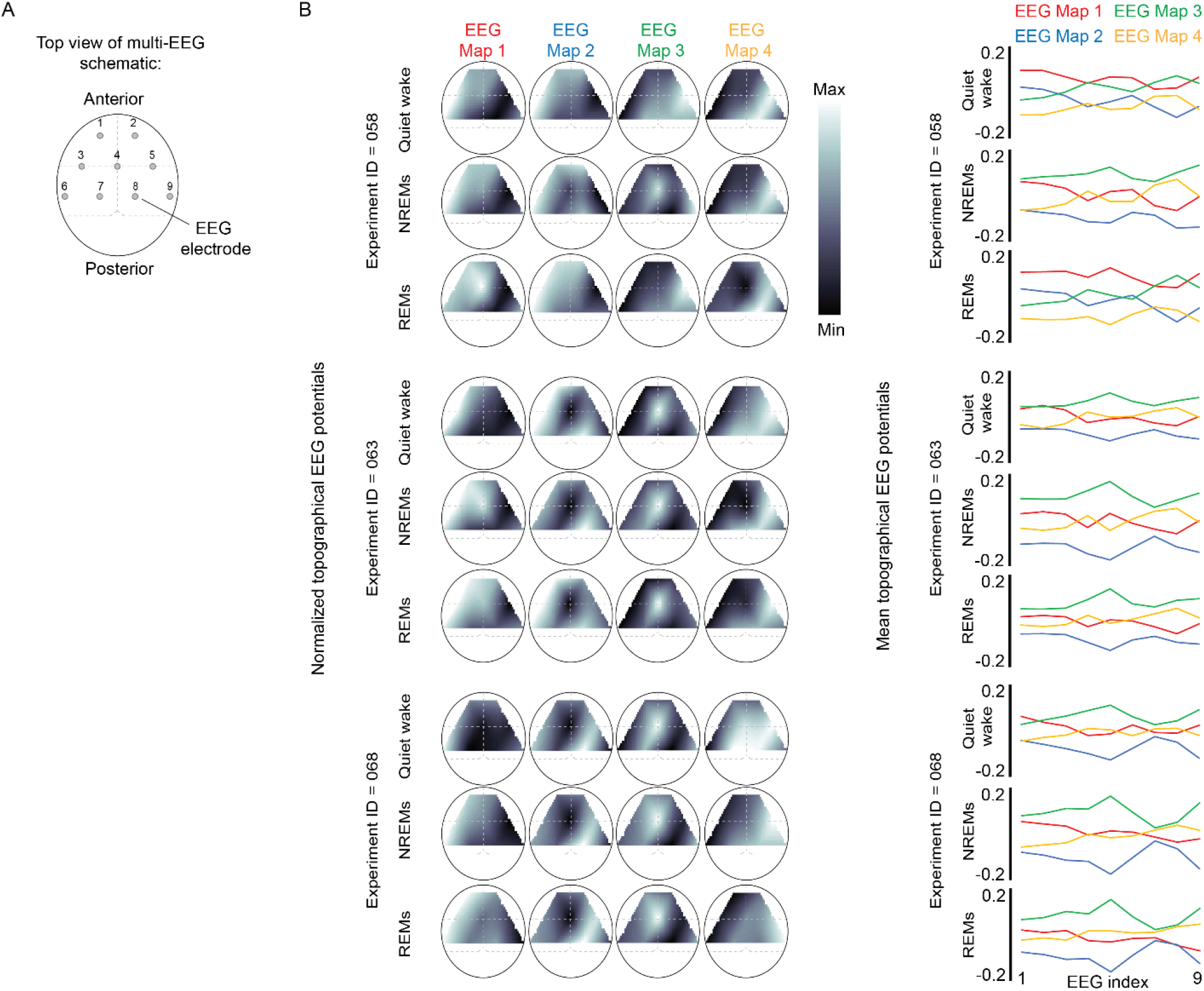
Comprehensive characterization of vigilance state-specific EEG microstates identified through k means clustering analysis of multi-EEG data. (**A**) Dorsal view schematic of multi-EEG configuration. Note that the imaging window is not included for clarity. (**B**) Normalized topographical EEG profile microstate maps derived from optimal significant cluster definitions for each vigilance state from each experiment (n=3 mice; experiment ID # 058, 063, 068). Data are scaled according to the max-min difference for each map individually. Right. Color-coded linear plots of the same topographical EEG maps to provide direct comparison of values for each map. For all figures in (**B**), EEG microstate maps have been ordered according to specific criteria (see supplementary methods).

**Fig. S3.**
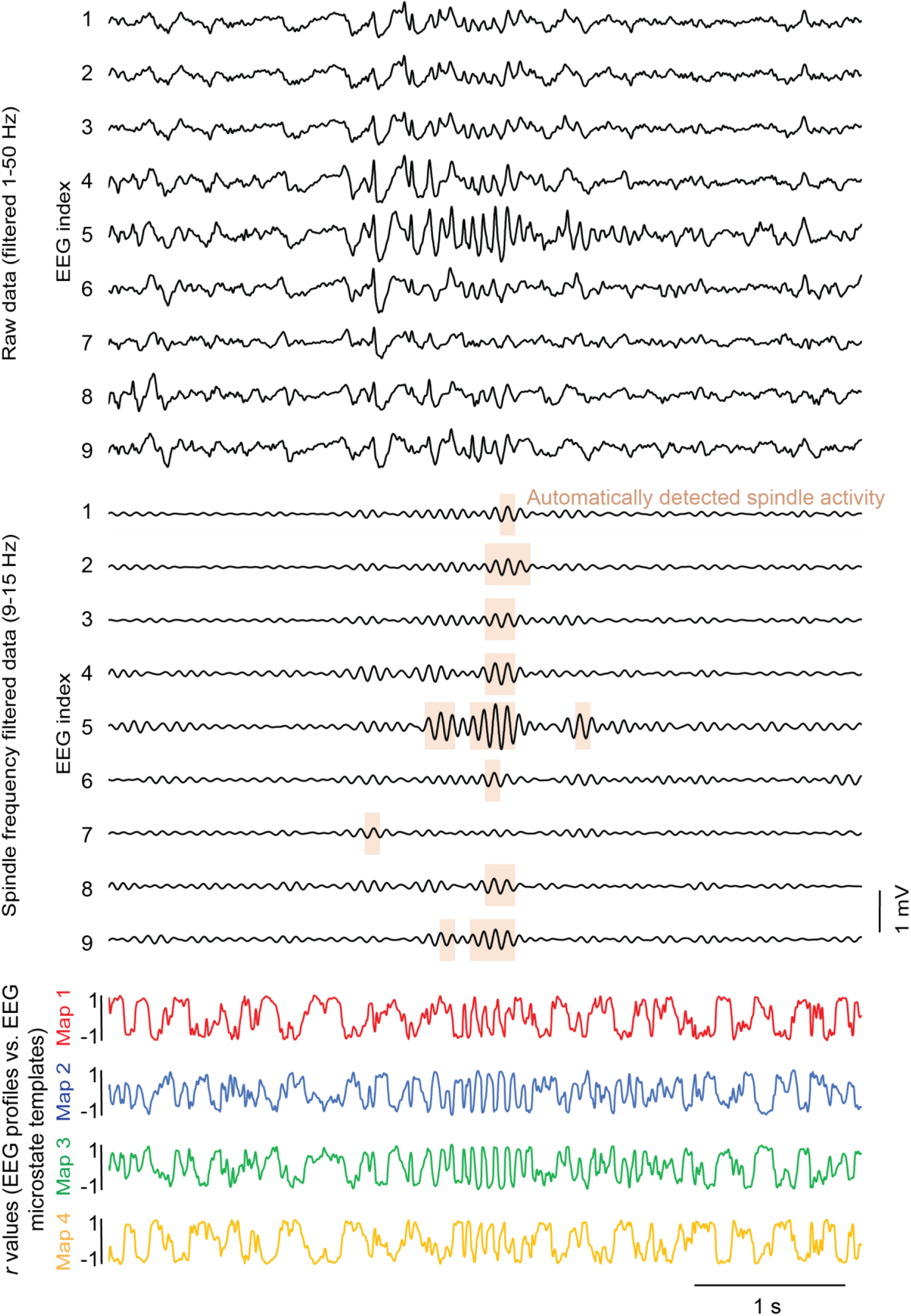
Automatic detection of significant spindle activity in multi-EEG recordings. Top. Example traces obtained from select NREMs data from one experiment. Middle. Spindle frequency (9-15 Hz) filtered version of above traces. Periods of significant spindle activity calculated (see supplementary methods for details on spindle detection procedure) for each EEG channel are highlighted in pink. Bottom. Plots of the corresponding Pearson correlation ***r*** values calculated between momentary multi-EEG topographical profiles (i.e., traces at top) and microstate map templates calculated for current select NREMs data at each time point.

**Fig. S4.**
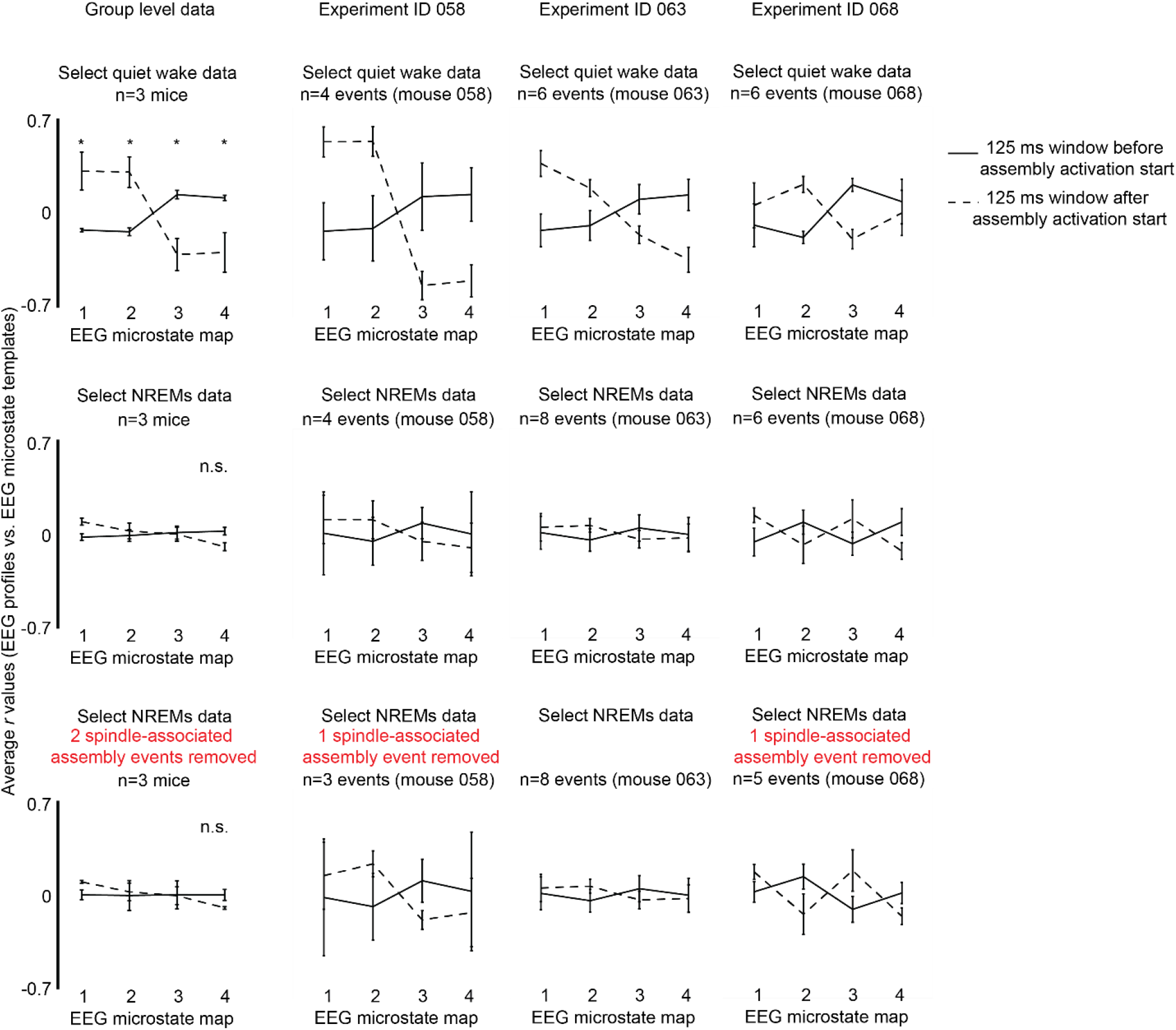
Comprehensive analysis of coordination between assembly activations in the sensorimotor cortex and EEG microstate dynamics. Top. At left, group level analysis of Pearson correlation ***r*** values calculated between momentary multi-EEG topographical profiles and microstate map templates at each data point within the +/- 125 ms windows surrounding assembly activation start points for quiet wakefulness. Analysis for each individual experiment during quiet wakefulness are included at right. Middle. Same as above, but data is for NREMs. Bottom. Same data as NREMs plots at middle, except spindle-associated events have been removed. (n.s.=not significant, \**P*<0.05, two-way repeated measures ANOVA with Holm-Šídák’s multiple comparisons post-hoc test, performed on group level data only). All group-level data are presented as mean ± SEM.

**Table S1.**
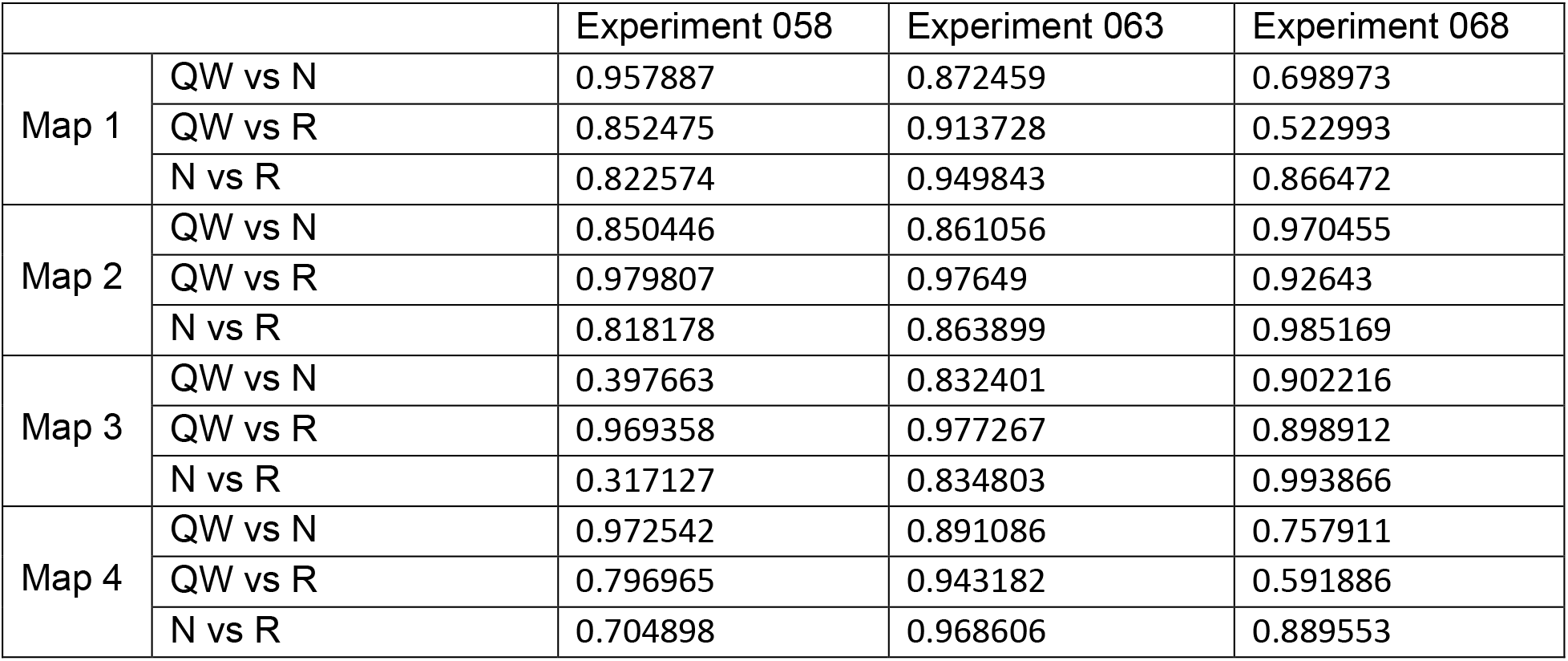
Comparison of EEG microstate map template similarity between different vigilance states within each experiment. Pearson correlation coefficient (***r***) values calculated between EEG microstate map templates derived from different vigilance states within the same multi-EEG experiment (n=3; experiment ID # 058, 063, 068).

|  |  | Experiment 058 | Experiment 063 | Experiment 068 |
| --- | --- | --- | --- | --- |
| Map 1 | QW vs N | 0.957887 | 0.872459 | 0.698973 |
|  | QW vs R | 0.852475 | 0.913728 | 0.522993 |
|  | N vs R | 0.822574 | 0.949843 | 0.866472 |
| Map 2 | QW vs N | 0.850446 | 0.861056 | 0.970455 |
|  | QW vs R | 0.979807 | 0.97649 | 0.92643 |
|  | N vs R | 0.818178 | 0.863899 | 0.985169 |
| Map 3 | QW vs N | 0.397663 | 0.832401 | 0.902216 |
|  | QW vs R | 0.969358 | 0.977267 | 0.898912 |
|  | N vs R | 0.317127 | 0.834803 | 0.993866 |
| Map 4 | QW vs N | 0.972542 | 0.891086 | 0.757911 |
|  | QW vs R | 0.796965 | 0.943182 | 0.591886 |
|  | N vs R | 0.704898 | 0.968606 | 0.889553 |

**Table S2.** Comparison of EEG microstate map template similarity for each vigilance state across experiments. Vigilance state-specific Pearson correlation coefficient (***r***) values calculated between EEG microstate map templates derived from different multi-EEG experiments (n=3; experiment ID # 058, 063, 068).

|  |  | Quiet wake | NREMs | REMs |
| --- | --- | --- | --- | --- |
| Map 1 | exp. 058 vs exp. 063 | 0.904675 | 0.91473 | 0.840512 |
|  | exp. 058 vs exp. 068 | 0.791621 | 0.929275 | 0.305193 |
|  | exp. 063 vs exp. 068 | 0.833449 | 0.856725 | 0.685786 |
| Map 2 | exp. 058 vs exp. 063 | 0.559141 | 0.569371 | 0.473702 |
|  | exp. 058 vs exp. 068 | 0.124694 | 0.376907 | -0.33357 |
|  | exp. 063 vs exp. 068 | 0.785267 | 0.880996 | 0.578107 |
| Map 3 | exp. 058 vs exp. 063 | 0.48335 | 0.739816 | 0.405163 |
|  | exp. 058 vs exp. 068 | 0.254769 | 0.755599 | -0.29438 |
|  | exp. 063 vs exp. 068 | 0.84878 | 0.901179 | 0.6585 |
| Map 4 | exp. 058 vs exp. 063 | 0.91437 | 0.926386 | 0.733376 |
|  | exp. 058 vs exp. 068 | 0.840129 | 0.931797 | 0.158764 |
|  | exp. 063 vs exp. 068 | 0.844393 | 0.854323 | 0.728752 |

**Table S3.** Relationship between EEG microstate map dynamics and individual EEG signals for each experiment. Pearson correlation coefficient (***r***) values calculated between EEG microstate map dynamics and 1-50 Hz bandpass filtered individual EEG recordings for each vigilance state from each multi-EEG experiment (n=3; experiment ID # 058, 063, 068).

|  |  |  | EEG 1 | EEG 2 | EEG 3 | EEG 4 | EEG 5 | EEG 6 | EEG 7 | EEG 8 | EEG 9 |
| --- | --- | --- | --- | --- | --- | --- | --- | --- | --- | --- | --- |
| Exp. 058 | Quiet wake | Map 1 | 0.72 | 0.66 | 0.35 | -0.26 | 0.13 | 0.32 | -0.51 | -0.54 | -0.01 |
|  |  | Map 2 | 0.63 | 0.55 | 0.25 | -0.37 | -0.04 | 0.22 | -0.55 | -0.66 | -0.19 |
|  |  | Map 3 | -0.64 | -0.55 | -0.25 | 0.37 | 0.04 | -0.22 | 0.55 | 0.65 | 0.19 |
|  |  | Map 4 | -0.72 | -0.66 | -0.36 | 0.24 | -0.14 | -0.34 | 0.50 | 0.56 | 0.00 |
|  | NREMs | Map 1 | 0.56 | 0.48 | 0.27 | -0.29 | 0.12 | 0.18 | -0.57 | -0.62 | -0.09 |
|  |  | Map 2 | 0.26 | 0.15 | 0.03 | -0.36 | -0.30 | -0.02 | -0.40 | -0.75 | -0.52 |
|  |  | Map 3 | 0.18 | 0.28 | 0.25 | 0.28 | 0.63 | 0.24 | 0.04 | 0.50 | 0.74 |
|  |  | Map 4 | -0.57 | -0.50 | -0.28 | 0.27 | -0.15 | -0.19 | 0.57 | 0.61 | 0.06 |
|  | REMs | Map 1 | 0.62 | 0.59 | 0.46 | -0.06 | 0.41 | 0.35 | -0.44 | -0.58 | 0.06 |
|  |  | Map 2 | 0.53 | 0.43 | 0.33 | -0.25 | 0.09 | 0.25 | -0.55 | -0.76 | -0.24 |
|  |  | Map 3 | -0.54 | -0.44 | -0.34 | 0.25 | -0.09 | -0.25 | 0.55 | 0.76 | 0.24 |
|  |  | Map 4 | -0.62 | -0.63 | -0.48 | -0.03 | -0.55 | -0.38 | 0.36 | 0.42 | -0.24 |
| Exp. 063 | Quiet wake | Map 1 | 0.40 | 0.55 | 0.34 | -0.48 | -0.20 | -0.19 | -0.56 | -0.59 | -0.14 |
|  |  | Map 2 | 0.02 | 0.11 | -0.05 | -0.59 | -0.68 | -0.38 | -0.46 | -0.63 | -0.56 |
|  |  | Map 3 | 0.02 | -0.07 | 0.09 | 0.57 | 0.74 | 0.37 | 0.41 | 0.56 | 0.53 |
|  |  | Map 4 | -0.42 | -0.57 | -0.35 | 0.46 | 0.14 | 0.18 | 0.56 | 0.59 | 0.13 |
|  | NREMs | Map 1 | 0.43 | 0.50 | 0.43 | -0.13 | 0.35 | 0.02 | -0.43 | -0.46 | 0.13 |
|  |  | Map 2 | -0.44 | -0.44 | -0.46 | -0.52 | -0.82 | -0.39 | -0.08 | -0.41 | -0.63 |
|  |  | Map 3 | 0.49 | 0.50 | 0.51 | 0.45 | 0.84 | 0.35 | 0.01 | 0.25 | 0.56 |
|  |  | Map 4 | -0.46 | -0.53 | -0.46 | 0.08 | -0.41 | -0.06 | 0.41 | 0.41 | -0.18 |
|  | REMs | Map 1 | 0.18 | 0.26 | 0.17 | -0.42 | -0.08 | -0.14 | -0.52 | -0.69 | -0.26 |
|  |  | Map 2 | -0.17 | -0.14 | -0.21 | -0.59 | -0.76 | -0.38 | -0.46 | -0.61 | -0.58 |
|  |  | Map 3 | 0.18 | 0.17 | 0.22 | 0.57 | 0.78 | 0.33 | 0.41 | 0.57 | 0.54 |
|  |  | Map 4 | -0.20 | -0.29 | -0.19 | 0.38 | 0.00 | 0.15 | 0.52 | 0.65 | 0.23 |
| Exp. 068 | Quiet wake | Map 1 | 0.69 | 0.30 | -0.01 | -0.51 | -0.38 | 0.06 | -0.52 | -0.43 | -0.05 |
|  |  | Map 2 | -0.18 | -0.56 | -0.70 | -0.86 | -0.91 | -0.69 | -0.68 | -0.68 | -0.80 |
|  |  | Map 3 | 0.04 | 0.46 | 0.65 | 0.87 | 0.90 | 0.61 | 0.70 | 0.67 | 0.72 |
|  |  | Map 4 | -0.75 | -0.40 | -0.10 | 0.41 | 0.27 | -0.16 | 0.46 | 0.38 | -0.05 |
|  | NREMs | Map 1 | 0.65 | 0.54 | 0.37 | -0.05 | 0.14 | 0.03 | -0.40 | -0.56 | -0.15 |
|  |  | Map 2 | -0.58 | -0.68 | -0.73 | -0.68 | -0.90 | -0.63 | -0.30 | -0.44 | -0.79 |
|  |  | Map 3 | 0.63 | 0.72 | 0.76 | 0.66 | 0.90 | 0.62 | 0.25 | 0.36 | 0.75 |
|  |  | Map 4 | -0.69 | -0.58 | -0.42 | 0.00 | -0.20 | -0.08 | 0.37 | 0.52 | 0.08 |
|  | REMs | Map 1 | 0.22 | 0.05 | 0.12 | -0.34 | -0.30 | -0.23 | -0.44 | -0.71 | -0.59 |
|  |  | Map 2 | -0.63 | -0.73 | -0.70 | -0.68 | -0.91 | -0.64 | -0.37 | -0.40 | -0.72 |
|  |  | Map 3 | 0.60 | 0.70 | 0.69 | 0.67 | 0.91 | 0.62 | 0.37 | 0.41 | 0.72 |
|  |  | Map 4 | -0.32 | -0.15 | -0.22 | 0.27 | 0.20 | 0.14 | 0.41 | 0.68 | 0.49 |

**Table S4.**
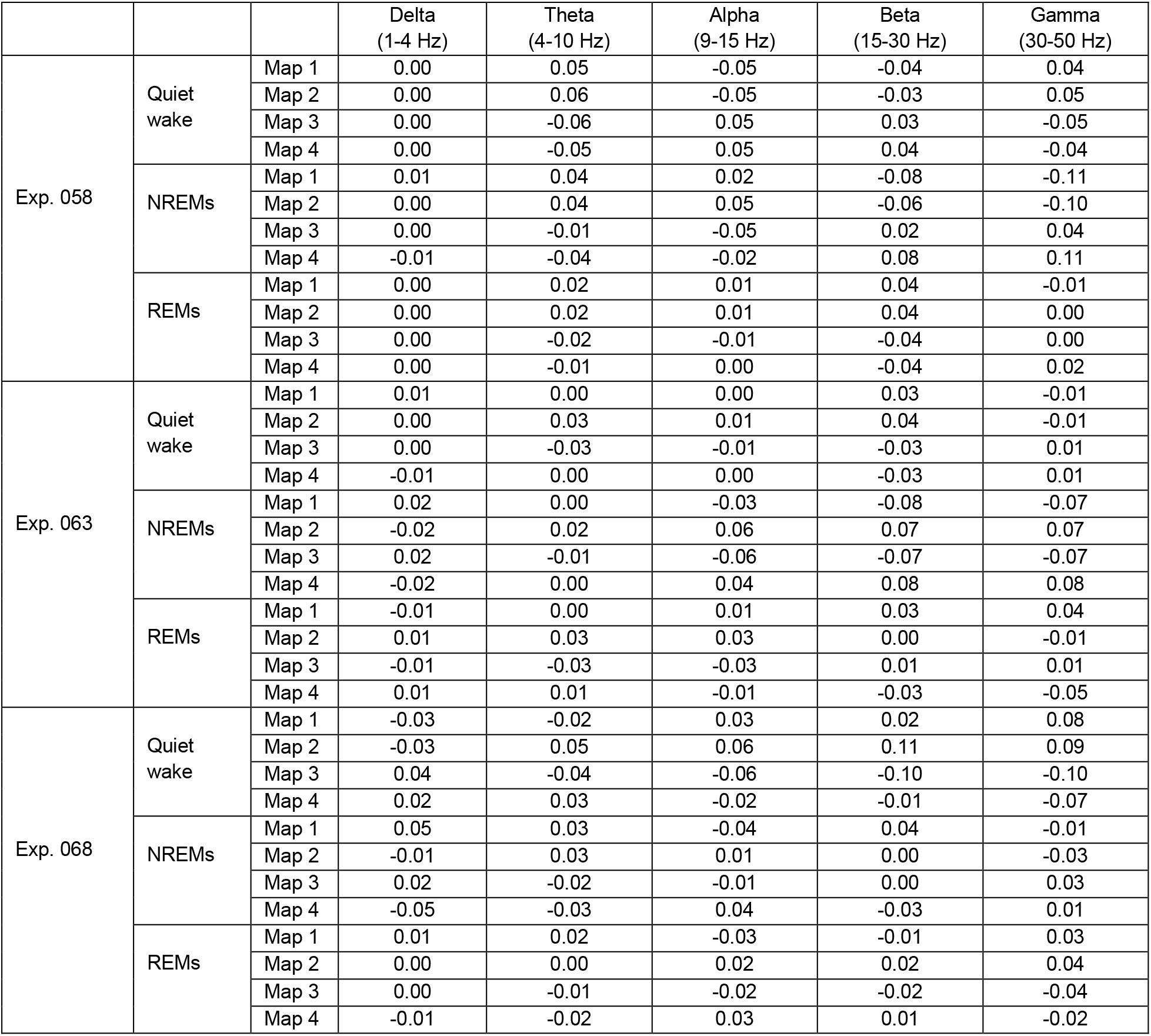
Relationship between EEG microstate map dynamics and global frequency band-specific EEG power for each experiment. Pearson correlation coefficient (r) values calculated between EEG microstate map dynamics and global wavelet transform analysis for each multi-EEG experiment individually (n=3; experiment ID # 058, 063, 068). For each frequency band tested (delta, theta, alpha, beta, gamma), the global wavelet transform value was calculated by first taking the sum of absolute wavelet amplitudes within the specified frequency range at each time point for each individual EEG. These values were then averaged across all EEG electrodes at each time point 0.5 s periods flanking temporal discontinuities in the select data were omitted from analysis.

